# Genomic analysis of fruit size and shape traits in apple: unveiling candidate genes through GWAS analysis

**DOI:** 10.1101/2023.08.21.554124

**Authors:** Christian Dujak, Maria José Aranzana

## Abstract

Genomic tools facilitate the efficient selection of improved genetic materials with within a breeding program. In this work, we focused on two apple fruit quality traits: shape and size. We utilized data from 11 fruit morphology parameters gathered across three years of harvest from 355 genotypes of the Apple REFPOP collection, which serves as a representative sample of the genetic variability present in European cultivated apples. The data was then employed for genome-wide association analysis (GWAS) using the FarmCPU and the BLINK models. The analysis identified 59 SNPs associated with fruit size and shape traits (35 with FarmCPU and 45 with BLINK) responsible for 71 QTNs. These QTNs were distributed across all chromosomes except for chromosome10 and 15. Thirty-four QTNs, identified by 27 SNPs, were related for size traits and thirty-seven QTNs, identified by 26 SNPs, were related to shape attributes. The definition of the haploblocks containing the most relevant SNPs served to propose candidate genes, among them the genes of the ovate family protein MdOFP17 and MdOFP4 which were in a 9.7kb haploblock on chromosome 11. RNA-seq data revealed low or null expression of these genes in the oblong cultivar ‘Skovfoged” and higher expression in the flat ‘Grand’mere’. In conclusion, this comprehensive GWAS analysis of the Apple REFPOP collection has revealed promising genetic markers and candidate genes associated with apple fruit shape and size attributes, providing valuable insights that could enhance the efficiency of future breeding programs.

## INTRODUCTION

Domesticated apples belong to the diploid species *Malus x domestica* (Suckow) Borkh. with a haploid chromosome number x = 17 and a highly duplicated genome of 651 Mb^1^. Parentage analysis performed in a large collection of European apple genotypes revealed a dense pedigree network with few key varieties highly used as founders at the top of the European pedigree, which remounts to few generations back^2,3^. Also, the contribution of the founders and their derived varieties to the overall pedigree was unequal.

Currently, a limited number of varieties dominate apple production and breeding, leading to a reduction in genetic diversity among commercial cultivars compared to that found in domesticated apples^4^. In breeding programs, fruit quality and productivity have traditionally been among the main objectives, while the recent need for varieties adapted to the effects of climate change (such as water scarcity, higher temperatures, and emerging diseases) demands more efficient and innovative breeding strategies, including novel phenotyping methods and molecular markers^5^. The use of molecular markers has enabled efficient selection in apple breeding, with different approaches being adopted in commercial breeding programs^6^. However, cutting edge scientific development and the availability of materials and genomic tools are essential for the progress of these breeding programs.

An essential tool for apple breeding is the REFPOP, a European collection of 534 genotypes (accessions and progenies) that represent the current European breeding germplasm. This collection was genotyped with high-density SNP arrays and evaluated over years in six European countries to study the environmental effect on the genotypes^7^. Using the REFPOP phenotypic and genotypic data, Jung et al.^8^ conducted genome-wide association (GWAS) and genomic selection (GS) studies, identifying important QTNs associated with numerous traits, including flowering time, harvested date, productivity, and fruit traits such as color, russeting, bitter pit, and fruit size, which need to be validated for use in breeding.

In addition to the above study, several works have aimed at identifying DNA polymorphisms associated with apple traits. Chagné et al.^9^ compiled a list of 128 single nucleotide polymorphisms (SNPs) for validation in a panel of accessions, including commercial varieties, advanced selections, and seedlings. Some of the SNPs were highly associated with relevant traits, making them suitable for molecular breeding.

Most of the works have been addressed to identify markers associated to disease resistance genes and to fruit quality traits like color, acidity, firmness, or compounds related to flavor. However, although few publications have focused on fruit size and shape, identifying genes or genomic regions that regulate these two important quality traits in apple and developing efficient markers for marker-assisted selection (MAS) is still challenging.

Significant progress has been made in understanding the genetic inheritance and regulation of fruit shape in vegetable crops. For instance, studies have identified several genes and QTLs that control the ovary and fruit elongation in tomato, such as SUN, OVATE, and FS8.1^10–12^. However, advances in apple fruit size and shape traits have been limited to the identification of a few molecular markers (SSRs and SNPs) located along the apple genome, except for chromosome 6^13–19^, but their efficiency for marker assisted selection is low. In addition, only few QTLs for fruit shape, measured as the ration between width and heigh (i.e. fruit shape index, FSI), have been identified in segregating populations^20–22^.

To enhance our understanding of the genomic regions, markers, and genes responsible for the inherited natural variation of fruit morphology, we conducted a GWAS study using fruit measurements obtained for a comprehensive description of fruit shape and size by^23^ in the densely genotyped apple REFPOP collection^7,8^. Also, whole genome RNA-seq data served to propose candidate genes that will require further validation.

## MATERIALS AND METHODS

### Plant materials

We used genotypic and phenotypic data of 355 genotypes of the Apple REFPOP copy growing in Gimenells (Lleida, Spain), including 257 accessions and 98 seedlings derived from 31 families (Supplementary Information 1).

### Genotypic data

Genotypic data were extracted from Jung et al.^8^ and consisted of 303,239 biallelic SNPs obtained with the Affymetrix Axiom^®^ Apple 480K SNP genotyping array^24^, or imputed from the Illumina Infinium^®^ 20K SNP genotyping array ^25^ in accessions and progenies, respectively.

### Phenotypic data

Phenotypic data were extracted from^23^, and consisted on four size and 10 morphometric descriptors obtained using the Tomato Analyzer software Version 3 developed by^26^ (Supplementary information 2). The data were collected in 12,692 apple sections harvested over three seasons: 2018 (134 genotypes), 2019 (274 genotypes) and 2020 (339 genotypes). Of these, 94 genotypes were evaluated in all three years. The descriptors included measures of size, fruit shape indexes, fruit blockiness, fruit homogeneity, distal fruit end shape, and internal fruit eccentricity. We also used the CAT-own fruit classification system to assign fruits into oblate or flat (class value = 1), spheroid or round (class value = 2), and oblong classes (class value = 3) based on visual comparison with images of three standard fruit typologies.

At least three apples per clone (two clones per genotype) and year were evaluated to obtain raw data. Mean values for each genotype were used for the analyses. For genotypes evaluated in more than one harvest season, mean values were calculated for each measure, resulting in a final dataset of 355 genotypes (referred as mean across-years dataset) (Supplementary Data 1).

Spearman’s correlation for all datasets, the distribution of the data and the density plots, and heatmaps were calculated and plotted with the *ggplot2* package^27^ in R Core Team (2022) program.

### Genome-wide association studies

Genome-wide association analysis (GWAS) were conducted using two methods. The Fixed and random model Circulating Probability Unification method (FarmCPU)^28^ and the Bayesian-information and Linkage-disequilibrium Iteratively Nested Keyway method (BLINK)^29^. FarmCPU combines the Mixed linear model with the Fixed Effect Model (FEM) to control for confounding factors, such as Kinship, and to reduce false negatives. It also incorporates the Random Effect Model (REM) to select associated markers by maximum likelihood method, thus avoiding the over-fitting. BLINK, on the other hand, replaces REM with FEM and uses the Bayesian information criteria (BIC) based on the linkage disequilibrium to generate fewer false positives and high statistical power.

Both FarmCPU and BLINK were implemented in the R package GAPIT 3.0^30^. GWAS was performed using genomic matrices with the same number of markers (303,239 SNPs) for four populations subsets with different sample sizes (as n_2018_=134 genotypes, n_2019_=274 genotypes, n_2020=_339 genotypes, and n_mean_=355). To control for population structure, we used three principal components. We filtered out SNPs with a minor allele frequency (MAF) < 0.05. To identify markers with significant association, we applied the Bonferroni correction with a significance threshold of a = *α/m*, where *α* = 0.05 and m is the number of markers (−*log*10(*p*-value) > 6.75).

The resulting p-values were plotted in multiple Manhattan and QQ plots using the threshold described^31^. Significant QTNs for all datasets and methods were graphically represented along each chromosome using the ggplot2 package^27^. Additionally, the GAPIT output file provided the phenotype variance explained by SNP (PVE) and the MAF. To further investigate the relationship between genotype and phenotype, we calculated the coefficient of determination using a numerical coding of alleles (1 and 2 for homozygous alleles and 3 for heterozygous alleles). The allelic frequency of each significant SNP was calculated with its corresponding association (phenotype), represented in boxplot using ggplot2.

### Haploblocks

We analyzed Linkage disequilibrium (LD) and identified Haploblocks using Haploview software^32^ based on the position of significant SNPs that were filtered using PLINK^33^. We focused on a 200kb window around the position of the SNP of interest (100kb each side), using the GDDH13 v1.1 genome^1^ as reference.

To identify haploblocks, we applied the following criteria: Hardy Weinberg p-value cut-off, 0.01; minimum genotype cut-off, 0.75; maximum number of Mendel errors, 1; minimum minor allele frequency, 0.05. We used the Gabriel et al.^34^ criteria to determine the blocks, which require a minimum confidence interval for strong LD (D’) at the top of 0.95 and at the bottom of 0.2 (indicating the LD level from 0.2 to 1).

Using Haploview software we calculated the allelic frequency of each haplotype in the population and identified connections between blocks.

### Candidate genes annotation

To annotate the genes in the haploblocks and the 200kb regions flanking the associated SNPs (100kb on both sides) we used the HFTH1 whole genome v1.0^35^ as the reference. For this, the haploblock regions initially aligned to the GDDH13 v1.1 whole genome assembly were subsequently aligned to the HFTH1 genome by BLAST+ from GDR database^36^. To further annotate these genes, we utilized various databases, including Gene Ontology (GO) terms^37^, InterPro (IPR)^38^, Kyoto Encyclopedia of genes and genomes (KEGG orthologs and pathways)^39^, non-redundant proteins sequences from NCBI (RefSeq)^40^, *Arabidopsis thaliana* orthologs from the Arabidopsis Information Resource (TAIR)^41^, and the computer-annotated protein sequence database for the translation of coding sequences (UniProtKB/TrEMBL) (The UniProt Consortium, 2019).

### RNA extraction and cDNA preparation

To investigate gene expression patterns, we collected apples from three genotypes with different shapes and sizes: ‘Grand’mere’ (GRA), flat and large size; ‘Kansas Queen’ (KAN), round and medium/small size; and ‘Skovfoged’ (SKO), oblong and medium size. Three biological replicates of fruit samples were collected at 13 days after anthesis. All fruit samples were frozen in liquid nitrogen and stored at −80°C until further processing. Total RNA was extracted from the frozen samples using the Maxwell® RSC simplyRNA tissue kit and the Maxwell® RSC instrument and was purify twice with Turbo® DNase to remove any residual DNA contamination. The quality and quantity of the extracted RNA was assessed using the Bioanalyzer system, and the RNA was set to Novogene London, England) for sequencing.

The RNA samples were converted to cDNA using the PrimeScript RT Reagent Takara kit. In the first step, the RNA was mixed with Oligo(dt) 20 nt (50 uM) and H_2_O RNase-free, and heated at 70°C for 5 min. In the second step, the cDNA synthesis reaction was performed using 5X PrimeScript Buffer, PrimeScript RT Enzyme, RNase out, dNTPs (100 uM), the first step reaction mixture, and H_2_O RNase-free, and incubated at 50°C for 60 min followed by inactivation at 70°C for 15 min. The resulting cDNA samples were then verified for integrity and subsequent user by visualizing them on a 1.5% agarose gel and 1X TAE.

### Analysis of mRNA sequencing data

The mRNA sequencing libraries were subjected to quality control, with reads having a Phred score < 30 being removed. Illumina sequencing adapters were trimmed using Trim-Galore^42^ version 0.6.1. Burrows-Wheeler Aligner ^43^ version 0.7.17 was used to map clean reads to the HFTH1 whole genome v1.High-quality RNA sequencing libraries were also mapped to HFTH1 by using HISAT^44^ version 2.1.0. with default settings parameters. SAMStat^45^ version 1.5.1 was used to analyze the quality of mapped and unmapped reads in Binary Alignment Map (BAM) files. The Mapping Quality Score (MAPQ) was used as an index to evaluate the quality of alignment and assembly, and only THE reads with MAPQ≥30 were retained. SAMtools^46^ version 1.9 was used to filter out multiple aligned reads and obtain statistic reports. Besides, filtered BAM files were transformed, indexed, and sorted according to the protocol needs using SAMtools. Quality control reports were generated before and after filtering and mapping with FastQC version 0.11.5^47^, and the results were summarized in an HTML file using MultiQC version 1.9^48^.

Gene quantification and count matrix construction were done with featureCounts^49^ applying paired-end sequencing parameters. Chimeric count fragments were avoided, and exon feature type was specified for read counting. The count matrix was annotated as transcript, and overlapping features were allowed for the differential use of exons during alternative splicing. The results were normalized to Transcript Per Million (TPM). To check for batch effects, the *sva* R package version 3.12^50^ was used. Preliminary exploratory analysis and visualization of the samples were also performed. The count matrix was normalized using a regularized logarithm transformation (*rlog*) to stabilize the variance across the mean for negative binomial data with a dispersion-mean trend and a low number of samples (n < 30).

The differential expression of genes (DEGs) annotated in the haploblock was analyzed using a Shapiro test to assess the normality of the distribution. For normally distributed data, an ANOVA-one way was applied, whereas a Kruskal-Wallis test was used for non-normal distribution. Differences between genotypes were determined the Tukey HSD test, with a confidence level of p<0.05, using the normalized count matrix in TPM.

### Validation RNA-seq and expressed genes (TPM)

We validated the differential expression of the gene HF43536 using the primers Fw 5’-AGGGCAGCTAAGGATTTGGA-3’ and Rv 5’-TGTGTGTGCCATGTCAAACCAG-3. The qPCR was performed using the LightCycler 480 System Roche. Each reaction contained 5x MasterMix SYBR Green, primers Fw and Rv (each 10uM), H_2_0 nuclease-free and cDNA adjusted to dilution 1:40. The cycling conditions were pre-incubation at 95°C during 5 min, for amplification 40 cycles (at 95°C to 10 sec, 60° C to 10 sec and 72°C to 30 sec), melting curve (at 95° to 5 sec, 65°C to 1 min) and finally cooling at 40°C to 1 min. The amplification efficiency was calculated using the formula Eff = −1 + 10^(−1/slope), and the RNAseq data was validated by analyzing the R-squared between log2(TPM) and the cycle threshold (ct).

### PhenoGram

A phenoGram^51^, based on chromosomal ideograms sharing the genomic information, was constructed using published SNPs or other molecular markers associated with apple shape and size. To locate them in the physical map the markers were first aligned using BLAST-NCBI^52^ with the double haploid GDHH13 v1.1 reference genome. All QTNs obtained in this study were also included.

## RESULTS

### Phenotypic data

The traits evaluated are broadly described in Dujak et al.^23^. For each trait, density plots were created to visualize how the data was distributed in each specific year. In addition, a density plot was generated to represent the mean distribution per trait across all years and give a sense of the overall trend or central tendency when combining the three years data (Supplementary Figure 1). Trait distributions tended to exhibit similarities in terms of their central tendencies and spread, suggesting that the traits being evaluated did not show significant changes or variations across the years and that mean values represented well the overall trend.

A comprehensive correlation analysis between traits, years, and the mean across years, revealed correlations ranging from moderate to strong (Figure 1, supplementary Figure 2). When considering the correlations between year and the mean across years values for a given attribute, the lowest value was found for the FST observations in 2019 (r=0.51) while the highest correlation was observed for FSII in 2020 (r=0.91) (Supplementary Figure 2).

**Figure 1.**
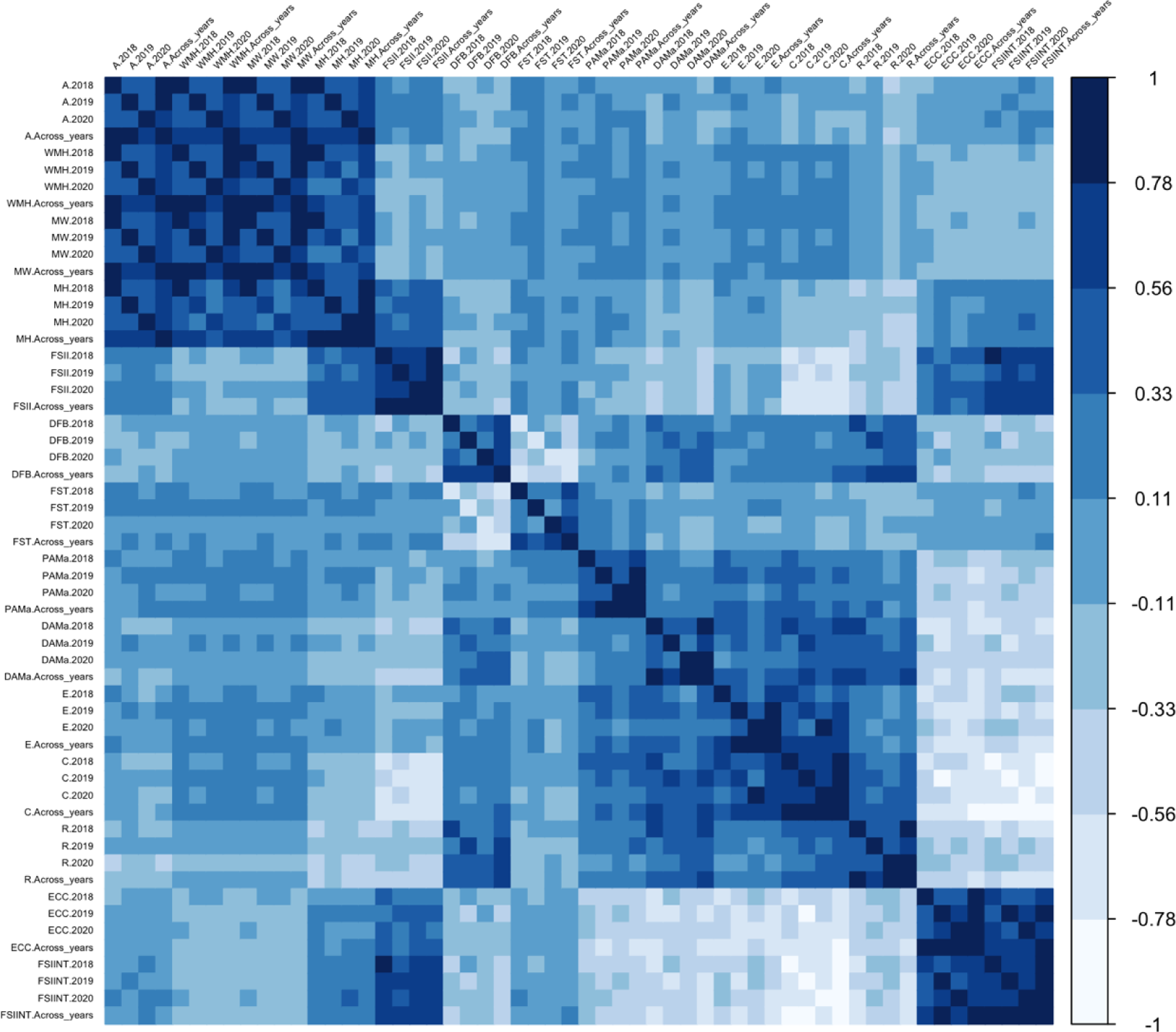
Spearman correlation analysis of fruit size and shape traits across multiple years, as well as their mean values across all years. The traits evaluated Area (A), Width Mid-height (WMH), Maximum Width (MW), Maximum Height (MH) Fruit shape index external I (FSII), Distal fruit blockiness (DFB), Fruit shape triangle (FST), Proximal angle macro (PAMa), Distal angle macro (DAMa), Ellipsoid (E), Circular (C), Rectangular (R), Eccentricity (ECC) and Fruit shape internal (FSIINT). See correlation coefficients in Supplementary Figure 2.

### Genome wide association studies

Genome-wide association studies (GWAS) were conducted for all traits using the per year as well as the mean across-years values using two models (FarmCPU and BLINK) (Figure 2). The results are displayed in Manhattan and QQ plots in the Supplementary Figures 3 and 4, respectively. The GWAS analysis identified SNPs with association values surpassing the Bonferroni threshold (−log10(p) =6.751) for all traits except for the fruit shape triangle (FST), for the distal angle macro (DAMa), ellipsoid (E) and for the eccentricity (ECC). Considering the two GWAS models, the three years of data and the mean across years values, we identified 59 SNPs associated (35 with FarmCPU and 45 with BLINK) responsible for 71 QTNs (Figure 2, Supplementary Table 1). Thirty-nine of the QTNs (55%) were identified when using the mean values. Five QTNs were identified simultaneously with two datasets (in all cases were QTNs detected with the 2020 and with the mean values datasets) and nine QTNs were identified by the two models in either one of the year’s assessments (six QTNs) or when using the means (three QTNs). In total, seven SNPs were simultaneously associated with more than one attribute, being one of the SNPs associated with three (AX-115482211 on chromosome 2, with A, MW and MWH). The 71 QTNs were distributed along all but the 10 and 15 chromosomes, ranging from two to 13 per chromosome. While some QTNs were scattered along the chromosome, others were in clusters.

**Figure 2.**
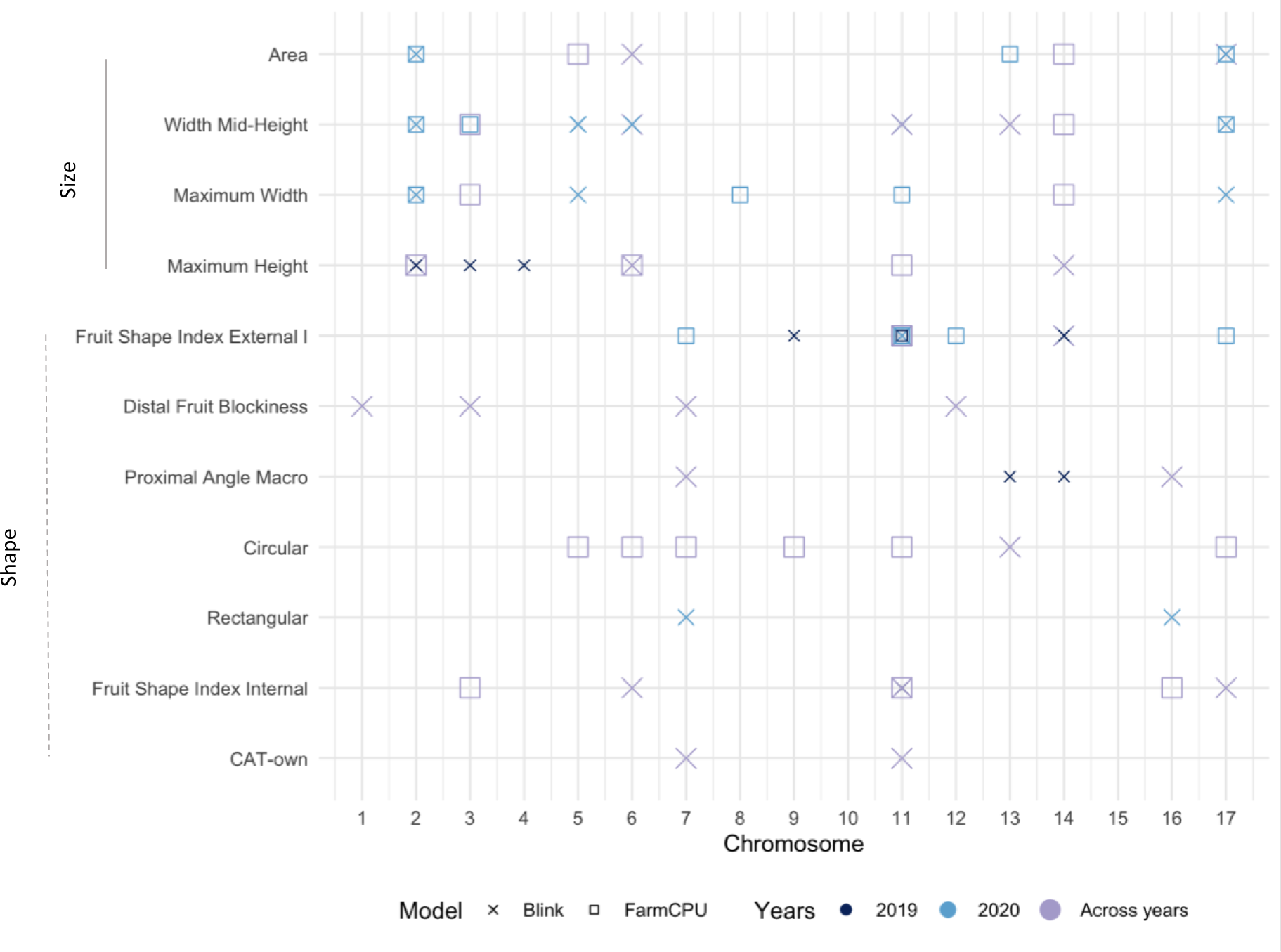
Summary of GWAS results for the size and shape traits using different models. QTNs obtained with Blink (x) and FarmCPU (≤) models in the datasets for 2019 (—), 2020 (—) and mean values (—) are represented. The X-axis corresponds to the chromosomes, while the Y-axis represents the traits.

### Quantitative Trait Nucleotides for size-related traits

Overall, our analysis revealed 34 QTNs significantly associated with size-related traits (Figure 2 and Supplementary Table 1). Among these, 12 QTNs were linked to width mid-height (WMH), eight to maximum height (MH), seven to maximum width (MW), and seven to area (A). A majority of the QTNs were discovered either in the dataset for the year 2020 or in the mean data encompassing all years. These QTNs were characterized by 27 SNPs, with five of these SNPs being associated with multiple QTNs. This is the case of the SNPs AX-115482211, on Chromosome 2, and AX-115481999 on Chromosome 3, identifying three QTNs each; and the SNPs AX-115378078 on Chromosome 6, AX-115295642 on Chromosome 14, and AX-115312607 on Chromosome 17, identifying two QNTs each.

Among the discovered SNPs, four exhibited simultaneous significance for both MW and WMH, localized on Chromosomes 2, 3, 14 and 17. The SNP on Chromosome 2 (AX-115482211) also demonstrated significance for the Area (A) trait (Supplementary Table 1). Moreover, the QTNs linked to WMH were distributed across eight different chromosomes.

### Quantitative Trait Nucleotides for shape-related traits

The study revealed 37 QTNs associated with shape-related traits, distributed across twelve chromosomes. Specifically, we identified twelve QTNs linked to the fruit shape index external I (FSII), seven associated with the circular measure (C), six with the fruit shape index internal (FSIINT), and four QTNs for each of the proximal angle macro and distal fruit blockiness measures. Additionally, we detected two QTNs each for the CAT-own and rectangular values.

Two significant SNPs on Chromosome 11 located approximately 32 Mb apart (AX-115335214 and AX-105213957) and one SNP on Chromosome 14 (AX-115336086) were responsible for six FSII QTNs for either the 2020 or the mean data. Furthermore, Chromosome 11 contained nine QTNs associated with C, FSII, FSIINT and CAT-own spaced along the chromosome; four QTNs (one for FSIINT, one for CAT-own and two for FSII) were in a region of 248kb (Supplementary Table 1).

Twenty-three of the QTNs were found when using the mean values, and 14 QTNs with the 2019 and 2020 datasets. Four out of the thirty-two SNPs were found simultaneously with the 2020 and the mean values datasets.

### Phenotypic variation and haploblocks

The phenotypic variation explained (PVE) by individual SNP spanned from 0.03% to 12.51%, with a mean PVE of 3.74 % (Supplementary Table 2). To visualize the genotype-phenotype relationship for each SNP and QTN, we used violin plots. Figure 3 illustrates 14 violin plots, each capturing the phenotypic differences among the three genotypic classes: homozygous for the reference allele, heterozygous, and homozygous for the alternative allele).

**Figure 3.**
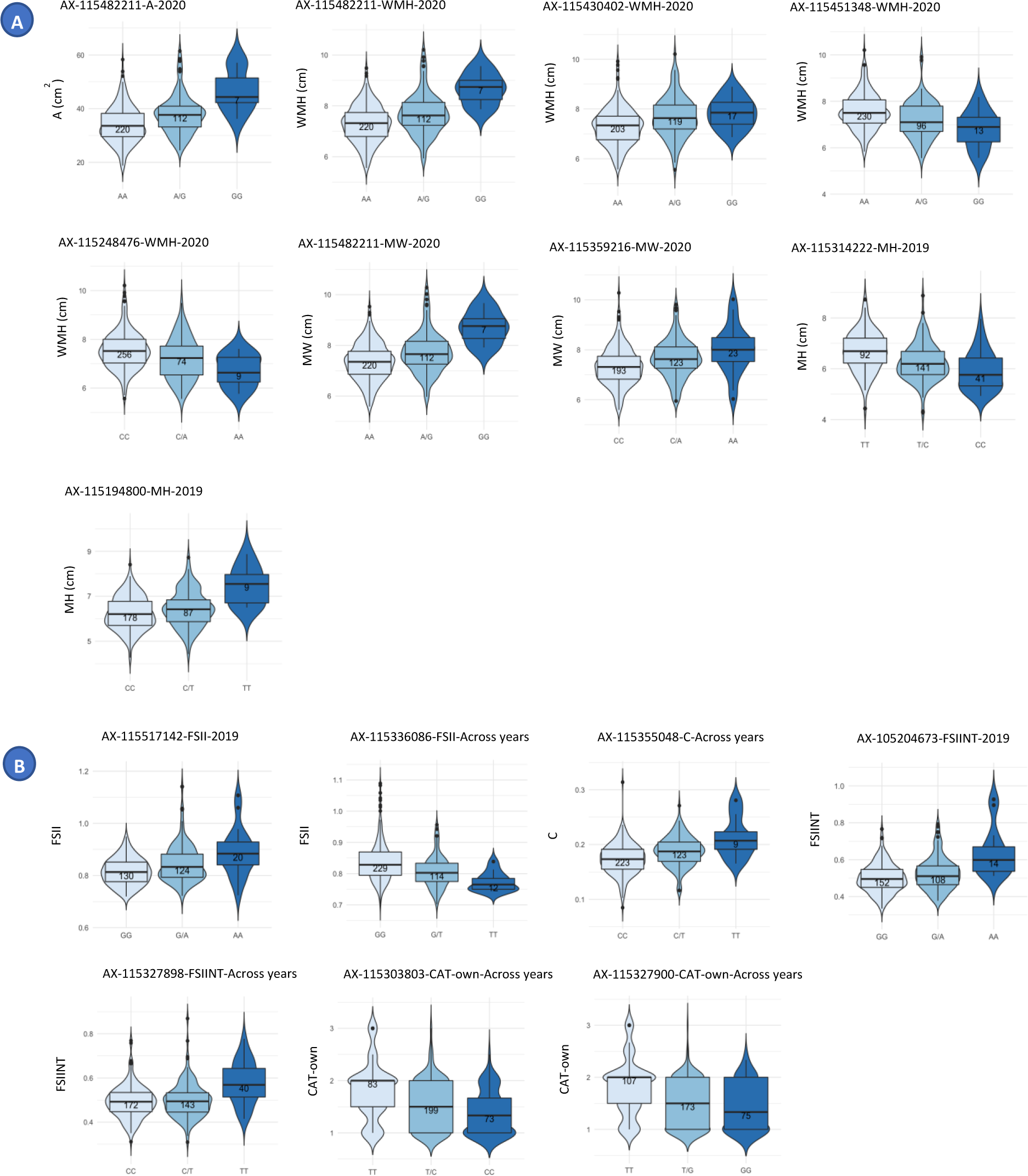
Violin plots displaying the frequency distribution of size (A) and shape (B) phenotypic values across genotypes. Each violin plot corresponds to a specific SNP-QTN combination, illustrating the distribution of trait values for different genotypes. In each violin plot, the X-axis represents the genotypes, with the first allele (on the left) indicating the homozygous genotype for the reference allele in the GDDH13 whole genome v1.1, the middle representing the heterozygous genotype, and the right side representing the homozygous genotype for the alternative or minor allele.

The SNP AX-115482211 displayed concurrent associations with three fruit size measures: Area, Width-mid height, and Maximum width. Individuals carrying the alternative allele G (with an allele frequency of 19%) exhibited significantly larger fruits compared to those with the reference allele. This effect was evident in both heterozygous and homozygous individuals. (Figure 3A and Supplementary Table 3).

To investigate the top 10 most outstanding SNP-QTN combinations, we conducted haplotype analysis by constructing haploblocks centered around the SNP. Due to linkage between two of the SNPs, we obtained a total of nine distinct haploblocks. These haploblocks were distributed in six chromosomes (2, 4, 6, 7, 11, and 13) and exhibited an average size of 31.5 kb, with lengths ranging from 1.1 to 111 kb. In total, 13 QTNs were found within these haploblocks (Supplementary Table 4).

A notable cluster of QTNs for both size and shape attributes occurred within a genomic region spanning 1.9 Mb along the chromosome 11. Among these QTNs, 11 were found within haploblocks, linked or co-segregating (Figure 4).

**Figure 4.**
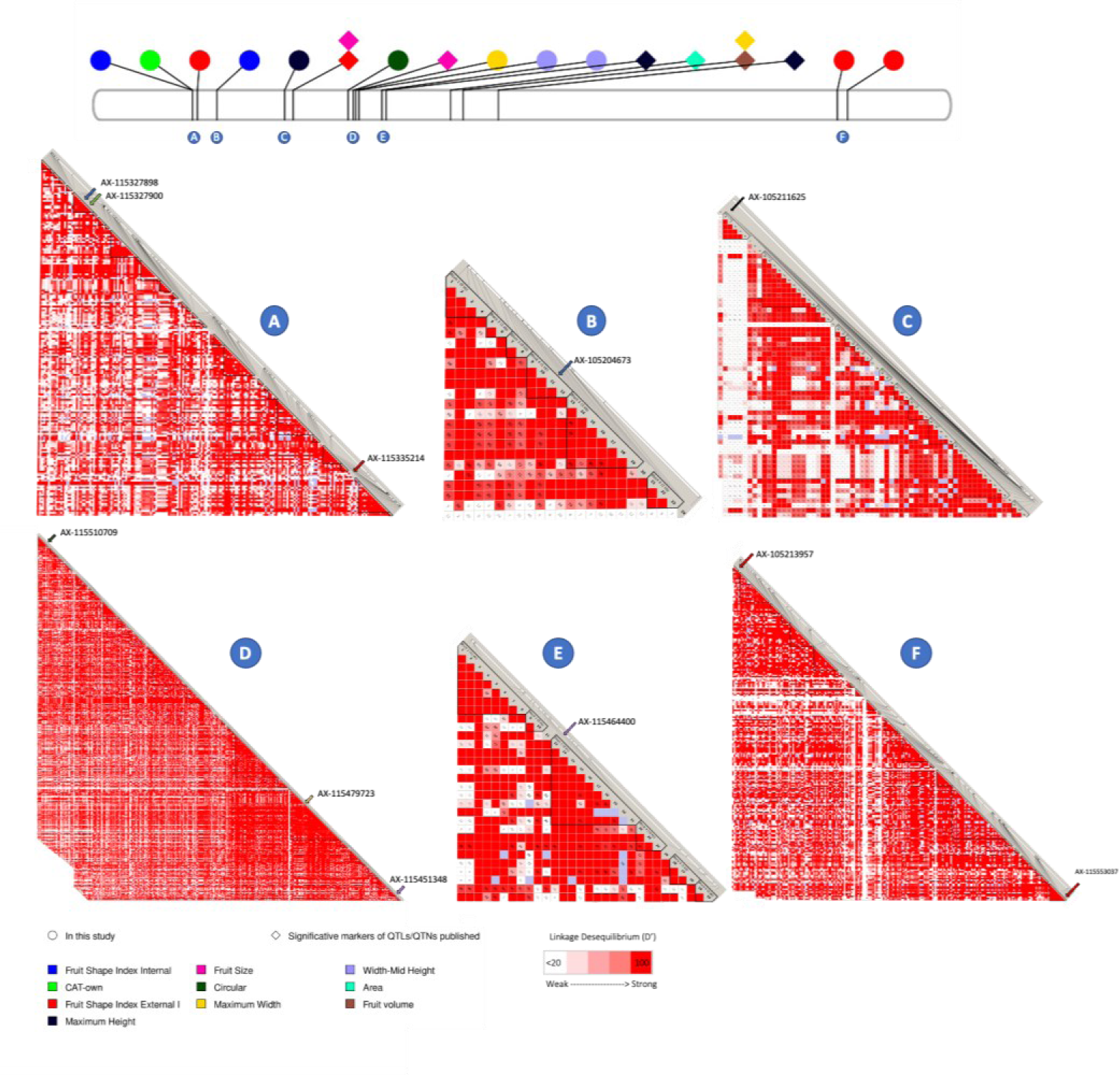
Linkage disequilibrium on Chromosome 11 GDDH13v1.1. The top figure displays Chromosome 11 from GDDH13v1.1, with position of markers published in this study and symbols representing QTNs found in this study (circles) and published (kites). The colors of the circles correspond to the associated QTN, and each circle is labeled with a letter representing the respective haploblock. Haploblocks, defined using GDDH13v1.1 positions, are as follows: Haploblock A: 4.661.534 to 4.958.286 (199 markers), Haploblock B: 5.930.883 to 5.948.789 pb (24 markers), Haploblock C: 9.046.670 to 9.556.662 pb (63 markers), Haploblock D: 12.638.696 to 13.198.304 pb (434 markers), Haploblock E: 14.355.224 to 14.402.233 pb (30 markers), Haploblock F: 37.648.782 to 38.174.120 bp (241 markers). The color gradient from white to red represents the level of linkage disequilibrium (D’). A D’ value <20 is considered weak linkage disequilibrium, and red color indicates a D’ value of 100, representing strong linkage disequilibrium.

In Figure 5A, we present three QTNs related to fruit shape, highlighting the respective haploblocks and their haplotype frequencies for three of the most significant associated SNPs: AX-115327898 (C/T alleles) and AX-115327900 (G/T alleles), both highly linked, situated 5kb apart at the top of chromosome 11 and associated to FSIINT and CAT-own attributes, respectively. Additionally, we identified AX-115355048 on chromosome 13 associated with the circular measure provided by the Tomato Analyzer software, which describes the extent to which the fruit section resembles a circle (Figure 5B).

**Figure 5.**
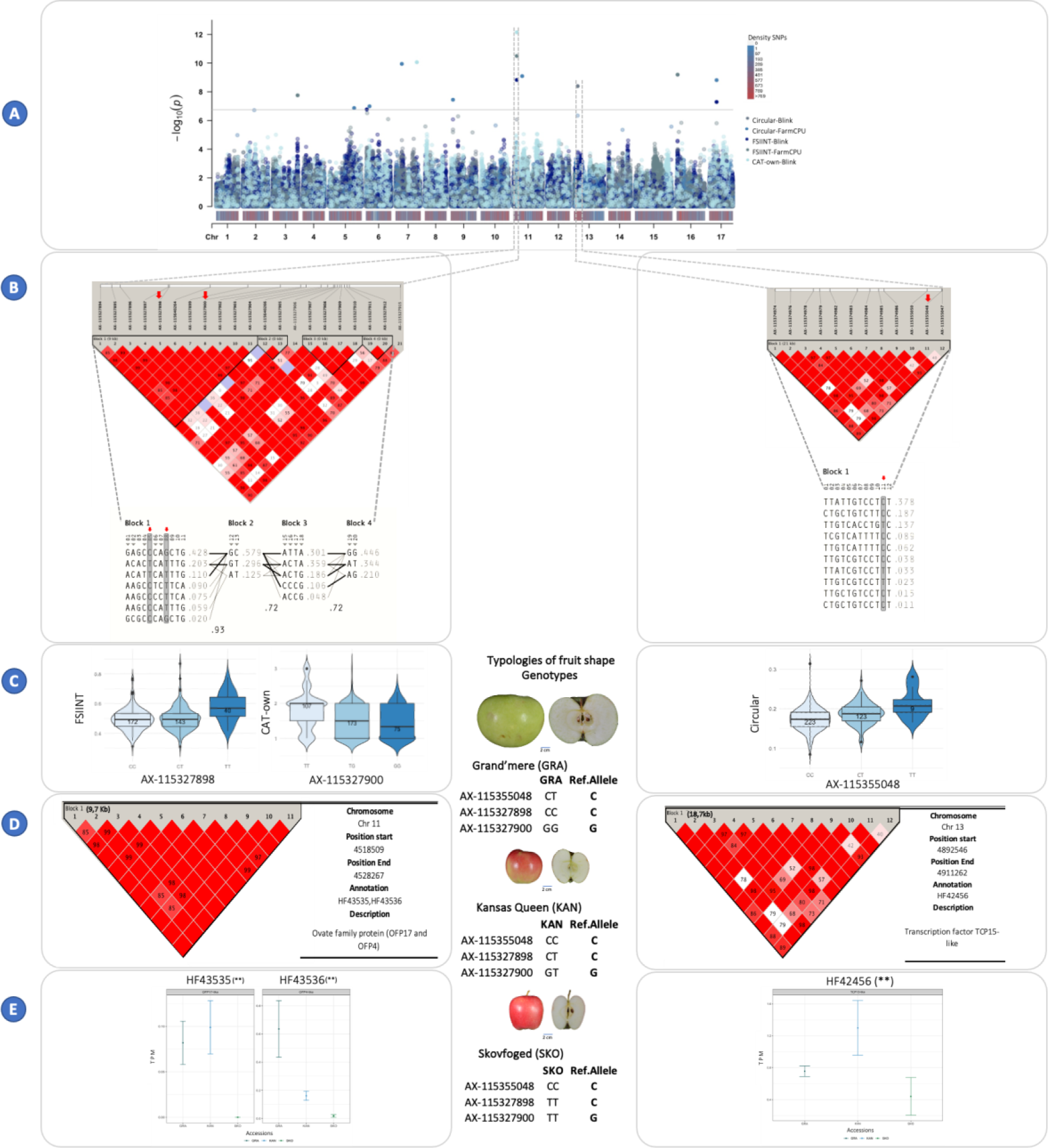
Integrated analysis of GWAS results, haploblocks, genotype-phenotype frequency, gene annotation, and RNA-seq, for FSIINT, CAT-own and Circular traits with the mean across years data. Panel **A, GWAS results:** this panel showcases multiple Manhattan plots representing the GWAS results for the three traits: FSIINT, CAT-own and Circular. The plots depict the significance of genetic markers on each chromosome, colored based on the trait and the two models used (Blink and FarmCPU). Density plots illustrate SNP distribution on each chromosome. Panel **B, Haploblocks & Haplotypes**: the linkage disequilibrium (D’) based on the GDDH13v1 genome is presented. Haploblocks are identified using the criteria from Gabriel et al. (2002), with colors indicating the strength of D’ (white = weak, red = strong). The haplotypes of each block and their allelic frequency are shown below. Panel **C, Frequency Genotype-Phenotype:** Allele frequency for the three traits is displayed, along with the corresponding apple shape genotypes (’Grand’mere’ = flat, ‘Kansas Queen’ = round, ‘Skovfoged’ = oblong). Panel **D, Gene annotation:** This section presents the candidate genes annotated within the haploblock, utilizing the annotations from the HFTH1 whole genome v1.0. Panel **E, RNA-seq:** Transcripts per million (TPM) data at the 13 Days After anthesis (DAA) fruit stage for three candidate genes (HF43535, HF43536 and HF42456) across the three genotypes.

Apples from cultivars with the allele T in AX-115327898 in homozygosis exhibited significantly higher FSIINT values, indicating a clear tendency for oblong fruit shapes. By contrary, individuals with CC and CT genotypes at this site produced flat and circular fruits (accessions such as ‘Grand’mere’ and ‘Kansas Queen’). Similarly, apples from cultivars homozygous for T in the SNP AX-115327900 such as ‘Skovfoged’ showed oblong shapes, while apples of heterozygous GT or homozygous GG cultivars were predominantly flat (as ‘Grand’mere’) or circular and ‘Kansas Queen’) (Figure 5C). These two SNPs were in complete LD (D’=1) and occurred in a haploblock 9,7 kb long, which had seven haplotypes with an average frequency of 0.14, ranging from 0.02 to 0.428 (Figure 5B).

The haploblock containing the SNP AX-115355048 on chromosome 13 (with CT alleles) was significantly associated with the circular attribute. The haploblock was 18,7 kb long and included 10 haplotypes with frequencies ranging from 0.378 to 0.011. Cultivars homozygous CC showed lower Circular values and higher FSIINT (see Figure 5C).

### Gene annotation

For each of the 59 associated SNPs, we searched for annotated genes within a 200 kb region (100 Kb upstream and downstream the SNP position) in the HFTH1 whole genome v1.0. This gene annotation analysis revealed 873 annotated genes, with 371 genes linked to size QTNs and 502 to genes linked shape QTNs. Additionally, the 53% of the annotations contained molecular description, according to Gene Ontology (GO) databases. Fifty-one genes had protein-binding molecular function, 40 genes were related to biological processes, including transcriptional regulation, DNA repair, phosphorylation, and transmembrane transport, among others. Moreover, we found genes that play vital roles in cell division, growth, cell modification and response to hormones, such as gibberellin, auxin, and ethylene.

Based on the TAIR database, a subset of genes was found to be directly linked to fruit development and growth. Specifically, we identified nine genes related to auxin response, including HF06172, HF40493, HF29276, HF02793, HF08237, HF41541, HF02644, HF02646, HF12008. Four genes were related to ethylene response (HF14170, HF14173, HF16534, HF11991), three genes involved in the gibberellins regulatory network (HF41950, HF38795, HF08230). Additionally, we recognized two genes related to fruit shape, including the Ovate Family protein HF43535 and HF43536 (Supplementary Table 5).

Gene annotation in the 9 haploblocks previously defined based on linkage disequilibrium, identified a total of 30 genes according to the TAIR database. Notably, we found among these the Ovate Family Proteins 17 (OFP17) and 4 (OFP14) (HF43535, HF43536), the TCP15-like transcription factor involved in plant regulation (HF42456), and several proteins of the kinase superfamily (Figure 5D, Supplementary Table 4, and Supplementary Table 5).

### RNA-seq gene expression (TPM)

Whole RNA sequence data of three genotypes, one oblate (‘Grand’mere’), one round (‘Kansas Queen’) and one oblong (‘Skovfoged’) obtained from fruits at 13 days after anthesis were analyzed to evaluate the expression in fruit of the 30 genes annotated in the haploblocks (Supplementary Table 5). Twenty-three of them were transcriptionally expressed in fruits of the three genotypes. A total of six genes exhibited differential expression between the genotypes: the genes HF43535 and HF43536 (OFP17 and OFP14, respectively) were annotated within the haploblocks of the SNPs AX-115327898 and AX-115327900 on Chromosome 11. The genes HF10079 and HF10080 (belonging to the Patched family and protein kinase proteins, respectively) were found in the haploblocks of the SNPs AX-115513701 and AX-115448691 SNPs on chromosome 6. The gene HF15994 (encoding an unknown function protein) was identified in the haploblock of the SNP AX-115194800 on chromosome 4, while the gene HF42456 (a transcription factor TCP15-like) was in the haploblock of the SNP AX-115355048 on chromosome 13. Among these genes, three have been described to play a crucial function in organ regulation and development: the OFP17, the OFP4, and the *TCP15-like gene*. For the OFP17, significant differences in expression were observed between the ‘Grand’mere’ (flat) and ‘Skovfoged’ (oblong) (GRAvsSKO) and between ‘Kansas Queen’ (round) and ‘Skovfoged’ (KANvsSKO). However, no significant differences in expression were found between ‘Grand’mere’ and ‘Kansas Queen’ (GRAvsKAN). This gene was expressed at a lower level in the oblong variety ‘Skovfoged’ (**Figure 5E and Supplementary Table 6**). The HF43536 gene (OFP4) was differentially expressed in the pairs GRAvsKAN (oblate and round) and GRAvsSKO (oblate and oblong), with higher RNA levels in the flat genotype.

As a mean to validate the RNA-seq data, the gene expression of this last gene (HF43536) was assessed by RT-qPCR, obtaining an Eff=2 and a r-squared of 0.8591 between the Cycle Threshold (Ct) and log2 (TPM) values (Supplementary Figure 5).

*Transcription factor TCP15-like gene (HF42456) showed d*ifferences in gene expression levels between the oblate and oblong fruits (GRAvsSKO) and between round and oblong fruits (KANvsSKO), with the SKO genotype showing lower gene expression (Figure 5E and Supplementary Table 6).

## DISCUSSION

Fruit shape, and in particular the shape of the apple, is relevant both for the description and varietal characterization as well as for aspects related to its commercialization and market value. Although visual and “easy to evaluate” criteria as the FSII for fruit classification is useful for the above-mentioned purposes, more objective data is necessary to perform genome studies. Here we used the data and measures obtained and described in^23^ to search for genomic regions controlling apple fruit shape and size attributes.

Fruit size and shape data was obtained in thousands of images acquired in fruits of a total of 355 genotypes in three consecutive harvesting campaigns (93 genotypes were common in the three years of assessments). It is broadly accepted that climatic and management factors affect fruit shape and size, although some studies show low differences in the FSII ratio between years, only observed under high divergences in the air and soil temperature in spring, for it may cause differences in the fruit seed number, the main factor determining fruit shape^53^. In the Spanish RefPop location, spring temperatures were only moderately milder in spring 2020, compared to 2018 and 2019, so we shouldn’t expect extreme divergences. The correlations between the values obtained each year support this fact.

GWAS analysis yielded significant associations for most of the studied traits, but there were a few exceptions where no significant associations were found. This finding might suggest that these traits have weaker genetic associations or that other factors beyond the examined SNPs play a more dominant role in influencing their expression or variation.

In our study, we made significant discoveries of several SNPs associated with size and shape-related traits. Joining these markers with markers and QTLs published in other studies^8,13,16,21,22^ we have constructed a PhenoGram with 110 molecular markers (SNPs and SSRs) (Figure 6, supplementary Table 7), 76 for size and 37 for shape traits.

**Figure 6.**
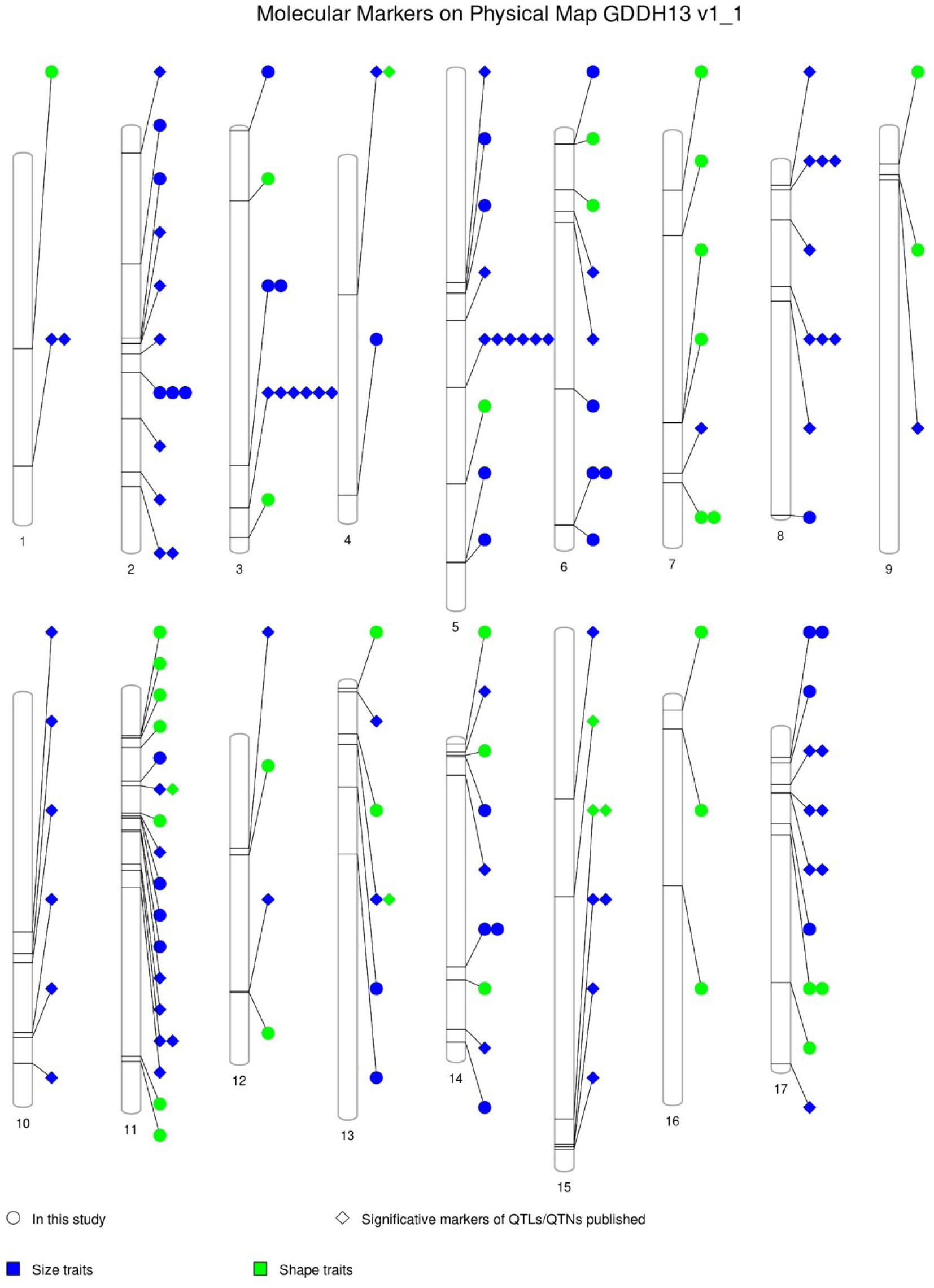
PhenoGram of the molecular markers on Physical map according to the apple GDDH13 whole genome v1.1. Significant markers mapped for apple fruit measures, including markers published in QTLs/QTNs analysis and the QTNs found in this study. Symbols: circle, correspond to “in this study” and kite, “significative markers of QTLs/QTNs published”. Color blue for Size traits and green for Shape traits. See details in Supplementary Figure 6.

### Size associated markers

Several size associated markers discovered here mapped close to other previously published. For example, on chromosome 3, ^16^ identified two QTLs responsible for fruit circumference and height in a segregating family. These QTLs collectively explained 45% of the phenotypic variation observed. Here we identified two QTNs (for MS and WMH) at 3.8Mb distance, with the gene HF40493 in the vicinity. This gene is a notable auxin response factor, sharing homology with *AtARF4* (identified by^54^), that plays a crucial role in regulating both female and male gametophyte development in Arabidopsis, as evidenced by the research conducted by^55^.

Also, on Chromosome 5, we found eight QTNs. Two of them (one for A and one for WMH) were at a very close distance (82 kb a part). The two associated SNPs (AX-115248476 and AX-115435503) were in LD (R-squared mean was 0.38) and added up to 10.6% of PVE. These QTNs were at about 15.7 Mb from a QTL for fruit maximum heigh reported by^16^ with a LOD of 3.94 and 21.5% of the variance explained. Two other QTNs for fruit width were at the top of this chromosome, with the two SNPs (AX-115638603 and AX-115436710) 102 kb apart and at less than 1Mb from a QTL for the same attribute identified by^21^ with a LOD of 2.9 explaining 9.2% of the variance, and 2.3 Mb apart from a QTL also for fruit width identified by^13^ with LOD3.5 and 12.4% of the variance explained. The SNPs explained together 6.57% of the variance. Similarly, ^16^ identified a width QTL 8.4Mb downstream of these SNPs.

Some genes annotated in these regions are responsible for growth regulation such as Transcriptional factor B3 family protein/auxin-responsive factor AUX/IAA-related (HF12008), ethylene responsive element binding factor 1 (HF11991), Gibberellin-regulated family protein (HF08230) and Auxin-responsive GH3 family protein (HF08237). Hormones, play an important role in fruit growth and are controlled by multiple genes. For example, endogenous auxin concentration is one of the factors controlling fruit size in apple^56^. In agreement with this, ^14^ suggest a potential role in fruit size of the Auxine Responsible Factor (ARF106) gene, contained in a QTL for fruit weight on Chromosome 15.

Chromosome 11 contains the highest number of markers (19) in the PhenoGram. For example, one of the associated SNPs here with Width mid-height (AX-115464400) is only 216 kb apart from the SNP AX-115380060 associated with fruit size in^8^. Also, the confidence interval of a fruit size-related QTL on this chromosome contains the miRNA172. The overexpression of this miRNA has a negative effect in fruit size^57^.

Additional QTLs/QTNs for fruit size attributes have also been reported along chromosomes 2, 8, 13, 14, and 17 in this study as well as in^8,13,14,21^.

### Shape associated markers

In this study, several QTNs for apple fruit shape attributes, measured through the analysis of bidimensional images, have been identified. Among the most relevant associations, we found a 9.7kb haploblock on Chromosome 11 with QTNs for FSIINT and CAT-own traits. The two associated SNPs (AX-115327898 and AX11532790) were in complete LD; the reference allele in homozygosis was present in the flat cultivars while the alternative in homozygosis was preferentially observed in oblong fruits. ^22^reported a QTL for the same measure (FSII), at 5 Mb distance from SNPs (AX-115327898 and AX-115327900) associated with FSII and another measure (CAT-own) that are highly correlated. ^21^ also detected several QTLs for fruit shape index (FSI), one of those QTLs in LG11 contributed to a phenotypic variance between 10.3-13.7% in a segregating population. This haploblock contained two ovate family protein genes (MdOFP17 and MdOFP4).

More than the 25% of the QTNs identified here are related to the fruit shape index (FSII and FSIINT studied here). This index has been preferentially used in successive works to describe fruit shape, since it is the measure with higher weigh in the definition of fruit shape^23^. However, other attributes are good descriptors of the apple shape, as is the fruit circularity (C). For this trait, the most relevant QTN was identified within an 18.7Kb haploblock on chromosome 13. The alternative allele of the associate SNP (AX-115355048) was preferentially observed in varieties bearing fruits with tendency to the oblong shape. This haploblock contained only one annotated gene, the TCP15-like transcription factor. This transcription factor is involved in the regulation of plant development and the stimulation of biosynthesis of hormones such as brassinosteroids, jasmonic acid, and flavonoids^58^.

### Genes expressed by RNA-seq data

#### Ovate family protein

The RNA-seq analysis of three varieties, each exhibiting contrasting phenotypes, revealed significant differences in their expression levels. For the gene MdOFP17, the oblong cultivar ‘Skovfoged’ showed lower expression levels, while the flat variety ‘Grand’mere’ showed an increase in the expression of MdOFP4. The two MdOFP genes occurred in cluster in the same haploblock.

The ovate family proteins are genes involved in the regulation of plant development in different organs, and in particular in the regulation of fleshy fruit shape, as described in several species such as tomato^59–61^, pepper^62^, melon^63^, and peach^64^. They are transcriptional repressor genes, but they also play an important role in the regulation of cell division in tomato fruit development, or in response to hormone changes^11,65^. In Arabidopsis, tomato and rice, the over-expression of OFPs causes the cotyledon, fruit, and seed to be flattened or, if there is a mutation in these genes, the organs are elongated ^66^ In apples, the diversity of OFP genes (26) distributed in 13 chromosomes has been studied^67^, but their role in apple fruit shape has not been described yet. Further investigation into the functions of these genes will shed light on their contributions to the observed differences in the studied plant populations.

## CONCLUSION

To date, few studies have been carried out to know which candidate regions or genes are responsible for fruit size and shape. In this study, we present QTNs and candidate genes for a better understanding of the genetic and molecular bases of apple fruit size and shape determination and highlight candidate genes, such as the MdOFP17 and MdOCP4, that may underlie the distinct fruit shapes observed among apple varieties. In addition, we provide here molecular markers for breeding.

## ACKNOWLEDGMENT

CD was supported by “DON CARLOS ANTONIO LOPEZ” Abroad Postgraduate Scholarship Program, BECAL-ParaguayThis research was supported by project PID2021-128885OB-I00 funded by MCIN/AEI/10.13039/501100011033 and by “ERDF A way of making Europe”. This project has received funding from the European Union’s Horizon 2020 research and innovation programme under grant agreement No 817970 (INVITE). We acknowledge support from the CERCA Programme (“Generalitat de Catalunya”), and the “Severo Ochoa Programme for Centres of Excellence in R&D” 2016-2019 (SEV-2015-0533) and 2020-2023 (CEX2019-000902-S) both funded by MCIN/AEI/10.13039/501100011033.

## AUTHOR CONTRIBUTION

MJA and CD contributed to the design and implementation of the research, to the analysis of the results and to the writing of the manuscript. CD conducted the laboratory experiments.

## REFERENCES

1 Daccord, N. et al. High-quality de novo assembly of the apple genome and methylome dynamics of early fruit development. Nat. Genet. 49, 1099 (2017).

2 Muranty, H. et al. Using whole-genome SNP data to reconstruct a large multi-generation pedigree in apple germplasm. BMC Plant Biol. 20, 1–18 (2020).

3 Pradas, N., Jurado-Ruiz, F., Onielfa, C., Arús, P. & Aranzana, M. J. PERSEUS: an interactive and intuitive pedigree visualization web application. Bioinformatics (submitted).

4 Gross, B. L., Henk, A. D., Richards, C. M., Fazio, G. & Volk, G. M. Genetic diversity in Malus×domestica (Rosaceae) through time in response to domestication. Am. J. Bot. 101, 1770–1779 (2014).

5 Laurens, F. et al. An integrated approach for increasing breeding efficiency in apple and peach in Europe. Hortic. Res. 5, 11, doi:10.1038/s41438-018-0016-3 (2018).

6 Migicovsky, Z. et al. Genomic consequences of apple improvement. Hortic. Res. 8 (2021).

7 Jung, M. et al. The apple REFPOP—a reference population for genomics-assisted breeding in apple. Hortic. Res. 7, 1–16, doi:10.1038/s41438-020-00408-8 (2020).

8 Jung, M. et al. Genetic architecture and genomic predictive ability of apple quantitative traits across environments. Hortic. Res. 9, doi:10.1093/hr/uhac028 (2022).

9 Chagné, D. et al. Validation of SNP markers for fruit quality and disease resistance loci in apple (*Malus x domestica* Borkh.) using the OpenArray® platform. Hortic. Res. 6, 30, doi:10.1038/s41438-018-0114-2 (2019).

10 Wu, S. et al. The control of tomato fruit elongation orchestrated by sun, ovate and fs8. 1 in a wild relative of tomato. Plant Sci. 238, 95-104 (2015).

11 Wang, S., Chang, Y. & Ellis, B. Overview of OVATE FAMILY PROTEINS, a novel class of plant-specific growth regulators. Front. Plant. Sci. 7, 417 (2016).

12 Mauxion, J.-P., Chevalier, C. & Gonzalez, N. Complex cellular and molecular events determining fruit size. Trends Plant Sci. 26, 1023–1038 (2021).

13 Kenis, K., Keulemans, J. & Davey, M. W. Identification and stability of QTLs for fruit quality traits in apple. Tree Genet. Genomes 4, 647–661 (2008).

14 Devoghalaere, F. et al. A genomics approach to understanding the role of auxin in apple (Malus x domestica) fruit size control. BMC Plant Biol. 12, 1–15 (2012).

15 Chagné, D. et al. Genetic and environmental control of fruit maturation, dry matter and firmness in apple (Malus× domestica Borkh.). Hortic. Res. 1 (2014).

16 Potts, S. M., Khan, M. A., Han, Y., Kushad, M. M. & Korban, S. S. Identification of quantitative trait loci (QTLs) for fruit quality traits in apple. Plant Mol. Biol. Rep. 32, 109–116 (2014).

17 Costa, F. MetaQTL analysis provides a compendium of genomic loci controlling fruit quality traits in apple. Tree Genet. Genomes 11, 819 (2015).

18 Sun, R. et al. A dense SNP genetic map constructed using restriction site-associated DNA sequencing enables detection of QTLs controlling apple fruit quality. BMC Genomics 16, 1–15 (2015).

19 Liu, Z. et al. Construction of a genetic linkage map and QTL analysis of fruit-related traits in an F1 Red Fuji x Hongrou apple hybrid. Open Life Sciences 11, 487–497 (2016).

20 Sun, H. H. et al. Identification of markers linked to major gene loci involved in determination of fruit shape index of apples (Malus domestica). Euphytica 185, 185–193 (2012).

21 Chang, Y. et al. Mapping of quantitative trait loci corroborates independent genetic control of apple size and shape. Sci. Hortic-amsterdam. 174, 126–132 (2014).

22 Cao, K. et al. Candidate gene prediction via quantitative trait locus analysis of fruit shape index traits in apple. Euphytica 206, 381–391 (2015).

23 Dujak, C., Jurado-Ruiz, F. & Aranzana, M. J. Comprehensive Morphometric Analysis of Apple Fruits and Weighted Class Assignation using Machine Learning Euphytica, doi:10.21203/rs.3.rs-2860631/v1 (*Under Review*).

24 Bianco, L. et al. Development and validation of the Axiom®Apple480K SNP genotyping array. Plant J. 86, 62–74, doi:10.1111/tpj.13145 (2016).

25 Bianco, L. et al. Development and validation of a 20K single nucleotide polymorphism (SNP) whole genome genotyping array for apple (*Malus x domestica* Borkh). Plos One 9, 9, doi:10.1371/journal.pone.0110377 (2014).

26 Gonzalo, M. J. et al. Tomato fruit shape analysis using morphometric and morphology attributes implemented in Tomato Analyzer software program. J. Am. Soc. Hortic. Sci. 134, 77–87 (2009).

27 Wickham, H. ggplot2: Elegant Graphics for Data Analysis. (Springer-Verlag New York, 2016).

28 Liu, X., Huang, M., Fan, B., Buckler, E. S. & Zhang, Z. Iterative usage of fixed and random effect models for powerful and efficient genome-wide association studies. Plos Genet. 12, e1005767 (2016).

29 Huang, M., Liu, X., Zhou, Y., Summers, R. M. & Zhang, Z. BLINK: a package for the next level of genome-wide association studies with both individuals and markers in the millions. Gigascience 8, giy154 (2019).

30 Wang, J. & Zhang, Z. GAPIT version 3: boosting power and accuracy for genomic association and prediction. Genomics, proteomics & bioinformatics 19, 629–640 (2021).

31 Yin, L. et al. rMVP: a memory-efficient, visualization-enhanced, and parallel-accelerated tool for genome-wide association study. Genomics, proteomics & bioinformatics 19, 619–628 (2021).

32 Barrett, J. C., Fry, B., Maller, J. & Daly, M. J. Haploview: analysis and visualization of LD and haplotype maps. Bioinformatics 21, 263–265 (2005).

33 Purcell, S. et al. PLINK: A tool set for whole-genome association and population-based linkage analyses. Am. J. Hum. Genet. 81, doi:10.1086/519795 (2007).

34 Gabriel, S. B. et al. The structure of haplotype blocks in the human genome. Science 296, 2225–2229 (2002).

35 Zhang, L. et al. A high-quality apple genome assembly reveals the association of a retrotransposon and red fruit colour. Nat. Commun. 10, 1–13 (2019).

36 Jung, S. et al. 15 years of GDR: New data and functionality in the Genome Database for Rosaceae. Nucleic Acids Res. 47, D1137–D1145 (2019).

37 Ashburner, M. et al. Gene Ontology: tool for the unification of biology. Nat. Genet. 25, 25–29, doi:10.1038/75556 (2000).

38 Hunter, S. et al. InterPro: the integrative protein signature database. Nucleic Acids Res. 37, D211–D215 (2009).

39 Kanehisa, M., Sato, Y., Kawashima, M., Furumichi, M. & Tanabe, M. KEGG as a reference resource for gene and protein annotation. Nucleic Acids Res. 44, D457–D462 (2016).

40 Pruitt, K. D., Tatusova, T. & Maglott, D. R. NCBI reference sequences (RefSeq): a curated non-redundant sequence database of genomes, transcripts and proteins. Nucleic Acids Res. 35, D61–D65 (2007).

41 Lamesch, P. et al. The Arabidopsis Information Resource (TAIR): improved gene annotation and new tools. Nucleic Acids Res. 40, D1202–D1210 (2012).

42 Krueger, F. & Andrews, S. R. Quality control, trimming and alignment of Bisulfite-Seq data (Prot 57). Department of Medicine, Hematology and Oncology, Domagkstr 3, 1–13 (2012).

43 Li, H. & Durbin, R. Fast and accurate long-read alignment with Burrows–Wheeler transform. Bioinformatics 26, 589–595 (2010).

44 Kim, D., Langmead, B. & Salzberg, S. L. HISAT: a fast spliced aligner with low memory requirements. Nat. Methods 12, 357–360 (2015).

45 Lassmann, T., Hayashizaki, Y. & Daub, C. O. SAMStat: monitoring biases in next generation sequencing data. Bioinformatics 27, 130–131 (2011).

46 Li, H. et al. The sequence alignment/map format and SAMtools. Bioinformatics 25, 2078–2079 (2009).

47 FastQC: a quality control tool for high throughput sequence data (available online at: http://www.bioinformatics.babraham.ac.uk/projects/fastqc, 2010).

48 Ewels, P., Magnusson, M., Lundin, S. & Käller, M. MultiQC: summarize analysis results for multiple tools and samples in a single report. Bioinformatics 32, 3047–3048 (2016).

49 Liao, Y., Smyth, G. K. & Shi, W. featureCounts: an efficient general purpose program for assigning sequence reads to genomic features. Bioinformatics 30, 923–930 (2014).

50 Leek, J. T., Johnson, W. E., Parker, H. S., Jaffe, A. E. & Storey, J. D. The sva package for removing batch effects and other unwanted variation in high-throughput experiments. Bioinformatics 28, 882–883 (2012).

51 Wolfe, D., Dudek, S., Ritchie, M. D. & Pendergrass, S. A. Visualizing genomic information across chromosomes with PhenoGram. BioData mining 6, 1–12 (2013).

52 Madden, T. The BLAST sequence analysis tool. The NCBI handbook (2003).

53 Tromp, J. Fruit shape in apple under various controlled environment conditions. Sci. Hortic-amsterdam. 43, 109-115 (1990).

54 Wang, C.-K. et al. Genome-wide analysis of auxin response factor (ARF) genes and functional identification of MdARF2 reveals the involvement in the regulation of anthocyanin accumulation in apple. New Zeal. J. Crop Hort. 49, 78–91 (2021).

55 Liu, Z. et al. ARF2–ARF4 and ARF5 are essential for female and male gametophyte development in Arabidopsis. Plant Cell Physiol. 59, 179–189 (2018).

56 Bu, H. et al. Endogenous auxin content contributes to larger size of apple fruit. Front. Plant. Sci. 11, 592540 (2020).

57 Yao, J. L. et al. A micro RNA allele that emerged prior to apple domestication may underlie fruit size evolution. Plant J. 84, 417–427 (2015).

58 Li, S. The Arabidopsis thaliana TCP transcription factors: a broadening horizon beyond development. Plant signaling & behavior 10, e1044192 (2015).

59 Brewer, M. T., Moyseenko, J. B., Monforte, A. J. & van der Knaap, E. Morphological variation in tomato: a comprehensive study of quantitative trait loci controlling fruit shape and development. J. Exp. Bot. 58, 1339–1349 (2007).

60 Tanksley, S. D. The genetic, developmental, and molecular bases of fruit size and shape variation in tomato. The plant cell 16, S181–S189 (2004).

61 Rodriguez, G. R. et al. Distribution of SUN, OVATE, LC, and FAS in the tomato germplasm and the relationship to fruit shape diversity. Plant Physiol 156, 275–285 (2011).

62 Zygier, S. et al. QTLs mapping for fruit size and shape in chromosomes 2 and 4 in pepper and a comparison of the pepper QTL map with that of tomato. Theor. Appl. Genet. 111, 437–445 (2005).

63 Martínez-Martínez, C. et al. A cryptic variation in a member of the Ovate Family Proteins is underlying the melon fruit shape QTL fsqs8. 1. Theor. Appl. Genet., 1-17 (2021).

64 Zhou, H. et al. A 1.7-Mb chromosomal inversion downstream of a PpOFP1 gene is responsible for flat fruit shape in peach. Plant Biotechnol. J. 19, 192, doi: 10.1111/pbi.13455 (2021).

65 Snouffer, A., Kraus, C. & van der Knaap, E. The shape of things to come: ovate family proteins regulate plant organ shape. Curr. Opin. Plant Biol. 53, 98–105 (2020).

66 Li, Q. et al. Molecular and genetic regulations of fleshy fruit shape and lessons from Arabidopsis and rice. Hortic. Res., uhad108, doi:10.1093/hr/uhad108 (2023).

67 Li, H., Dong, Q., Zhao, Q. & Ran, K. Genome-wide identification, expression profiling, and protein-protein interaction properties of ovate family proteins in apple. Tree Genet. Genomes 15, 1–11 (2019).

