## Supplementary material for "Genomic analysis of fruit size and shape traits in apple: unveiling candidate genes through GWAS analysis": Supplementary_Figures.pdf

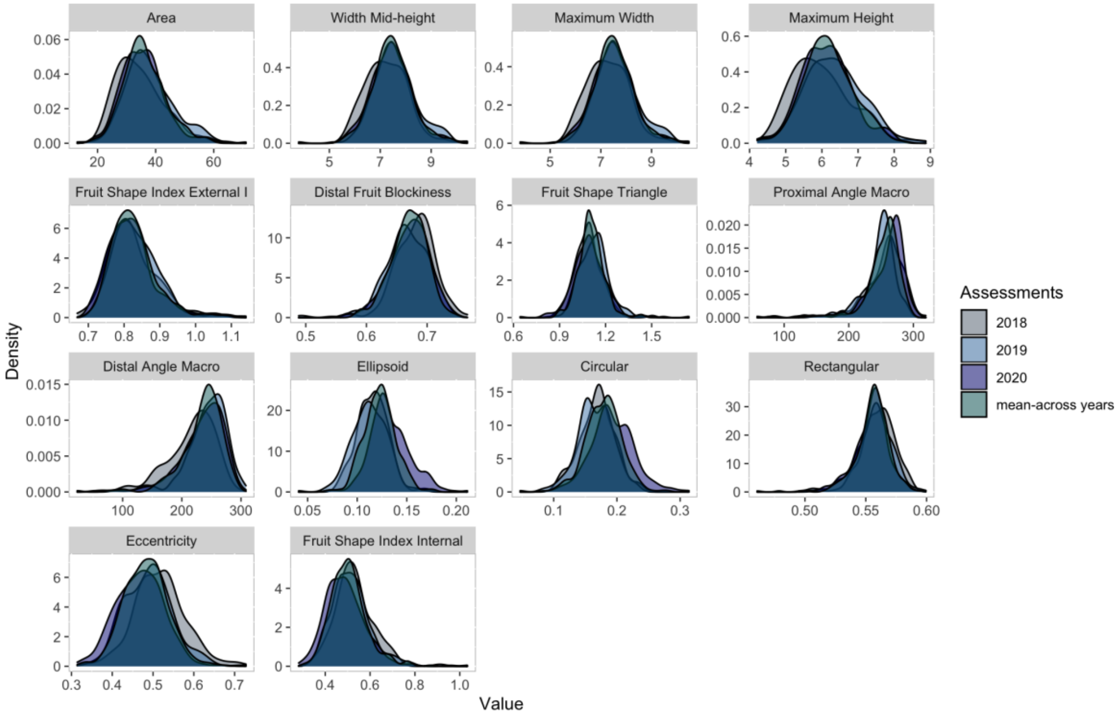


**Supplementary Figure 1**: Density plots illustrating distribution of the data for the three evaluated years (2018, 2019 and 2020) and the mean across all years, which gives a sense of the overall trend or central tendency across the years.

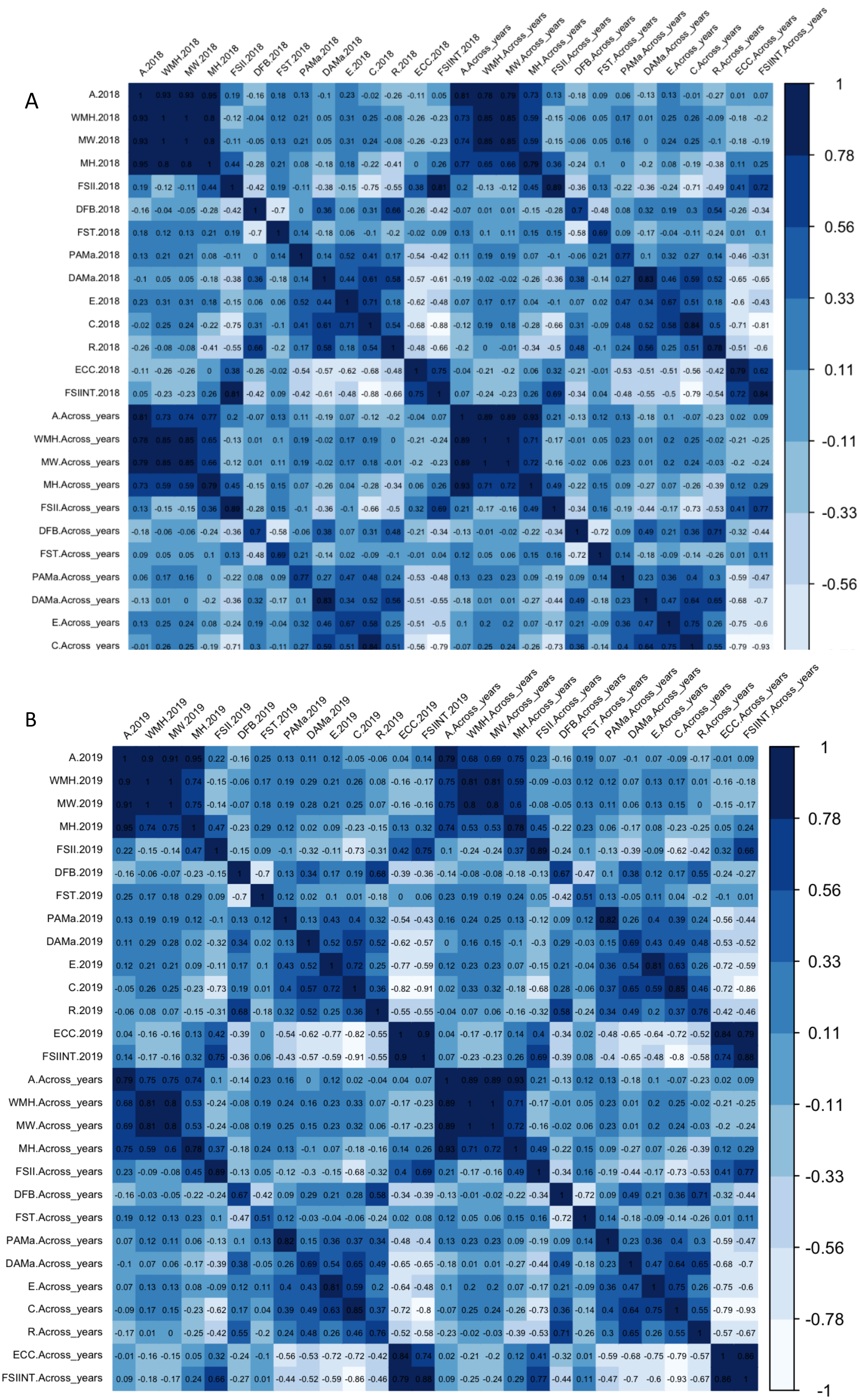
A
A.2018
WMH.2018
MW.2018
MH.2018
FSII.2018
DFB.2018
FST.2018
PAMa.2018
DAMa.2018
E.2018
C.2018
R.2018
ECC.2018
FSIINT.2018
A.Across_years
WMH.Across_years
MW.Across_years
MH.Across_years
FSII.Across_years
DFB.Across_years
FST.Across_years
PAMa.Across_years
DAMa.Across_years
E.Across_years
C.Across_years
R.Across_years
ECC.Across_years
FSIINT.Across_years
1
0.78
0.56
0.33
0.11
-0.11
-0.33
-0.56
B
A.2019
WMH.2019
MW.2019
MH.2019
FSII.2019
DFB.2019
FST.2019
PAMa.2019
DAMa.2019
E.2019
C.2019
R.2019
ECC.2019
FSIINT.2019
A.Across_years
WMH.Across_years
MW.Across_years
MH.Across_years
FSII.Across_years
DFB.Across_years
FST.Across_years
PAMa.Across_years
DAMa.Across_years
E.Across_years
C.Across_years
R.Across_years
ECC.Across_years
FSIINT.Across_years
1
0.78
0.56
0.33
0.11
-0.11
-0.33
-0.56
-0.78
-1

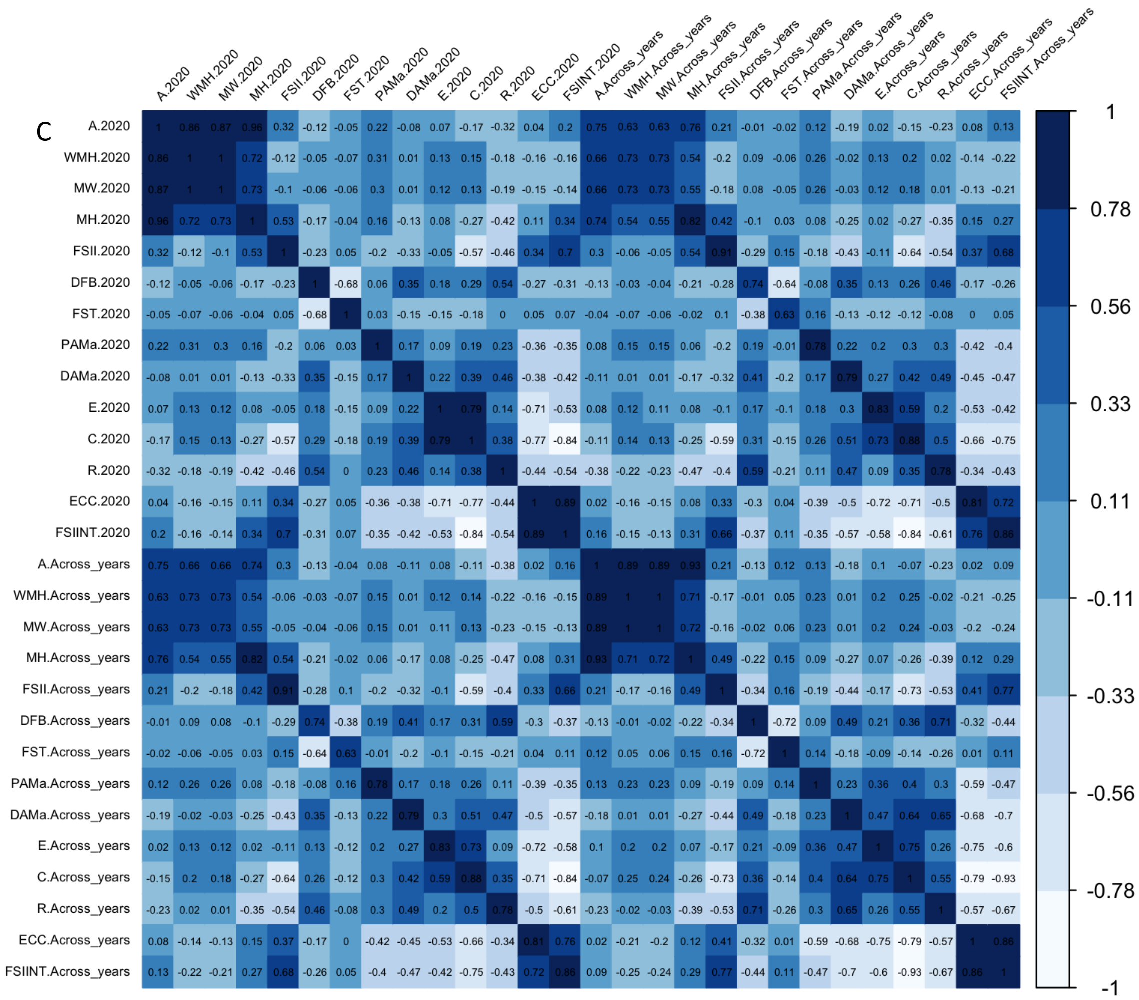


**Supplementary Figure 2**: Spearman correlation coefficients for each combination of traits in A) 2018, B) 2019, and C) 2020, including the mean across years values. The correlation scores are represented within individual squares. The traits evaluated include Area (A), Width Mid-height (WMH), Maximum Width (MW), Maximum Height (MH) Fruit shape index external I (FSII), Distal fruit blockiness (DFB), Fruit shape triangle (FST), Proximal angle macro (PAMa), Distal angle macro (DAMa), Ellipsoid (E), Circular (C), Rectangular (R), Eccentricity (ECC) and Fruit shape internal (FSIINT).

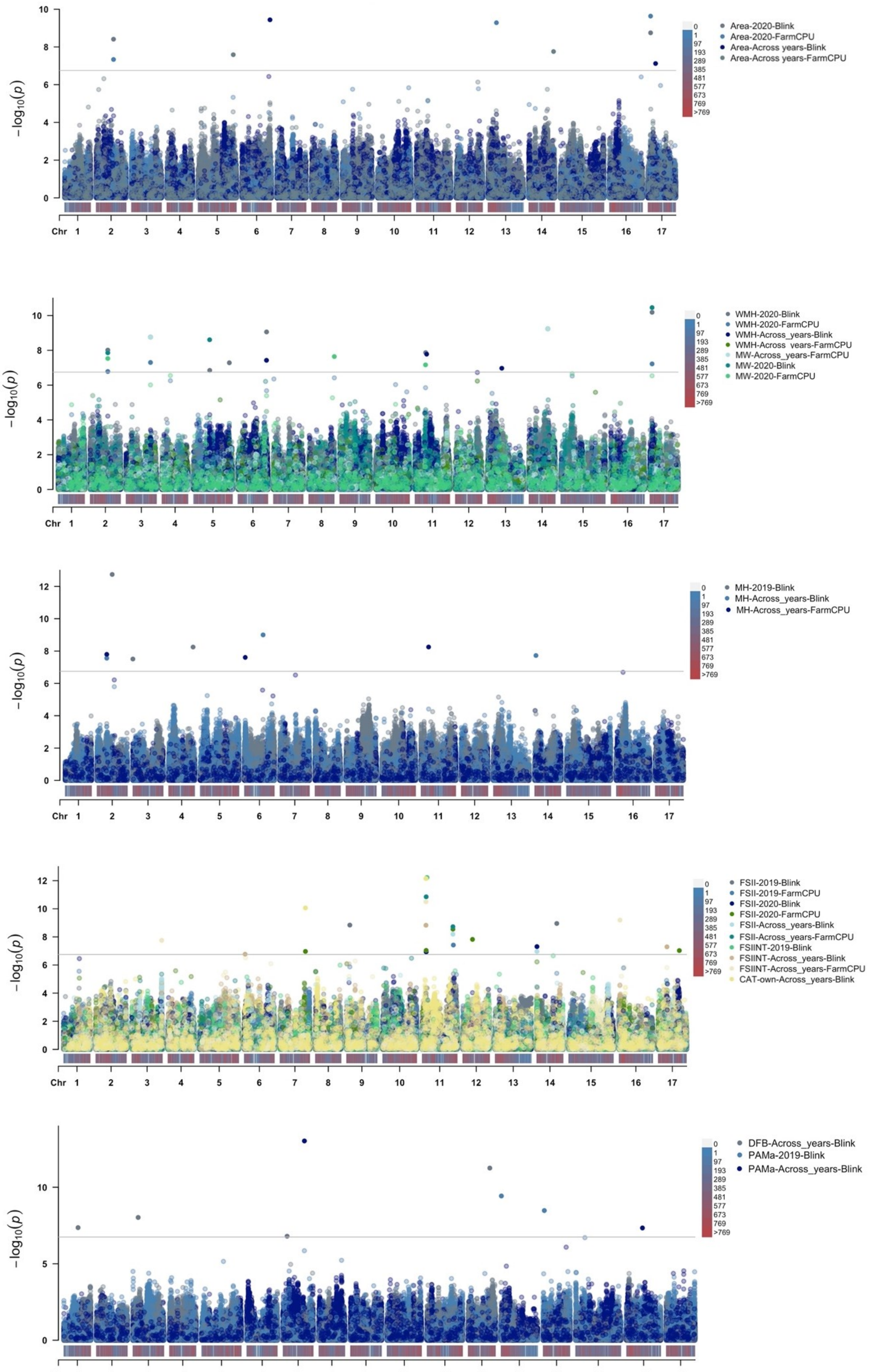
Area-2020-Blink
Area-2020-FarmCPU
Area-Across years-Blink
Area-Across years-FarmCPU
WMH-2020-Blink
WMH-2020-FarmCPU
WMH-Across_years-Blink
WMH-Across_years-FarmCPU
MW-Across_years-FarmCPU
MW-2020-Blink
MW-2020-FarmCPU
MH-2019-Blink
MH-Across_years-Blink
MH-Across_years-FarmCPU
FSII-2019-Blink
FSII-2019-FarmCPU
FSII-2020-Blink
FSII-2020-FarmCPU
FSII-Across_years-Blink
FSII-Across_years-FarmCPU
FSIINT-2019-Blink
FSIINT-Across_years-Blink
FSIINT-Across_years-FarmCPU
CAT-own-Across_years-Blink
DFB-Across_years-Blink
PAMa-2019-Blink
PAMa-Across_years-Blink
$-\log_{10}(p)$
Chr

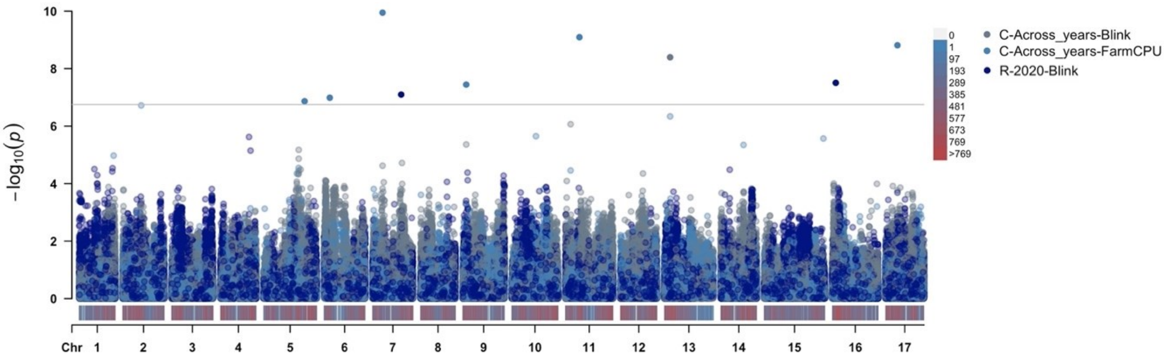


**Supplementary Figure 3**: Manhattan plots displaying GWAS results for each phenotype per year and across all years, along with the GWAS model used. The Bonferroni threshold (6,751) is shown as a gray line for reference. The density of SNPs per chromosome is represented by a color gradient on the X-axis, ranging from gray (lower density) to red (higher density). Each plot is labeled with the corresponding trait-year or across years model in the legend.

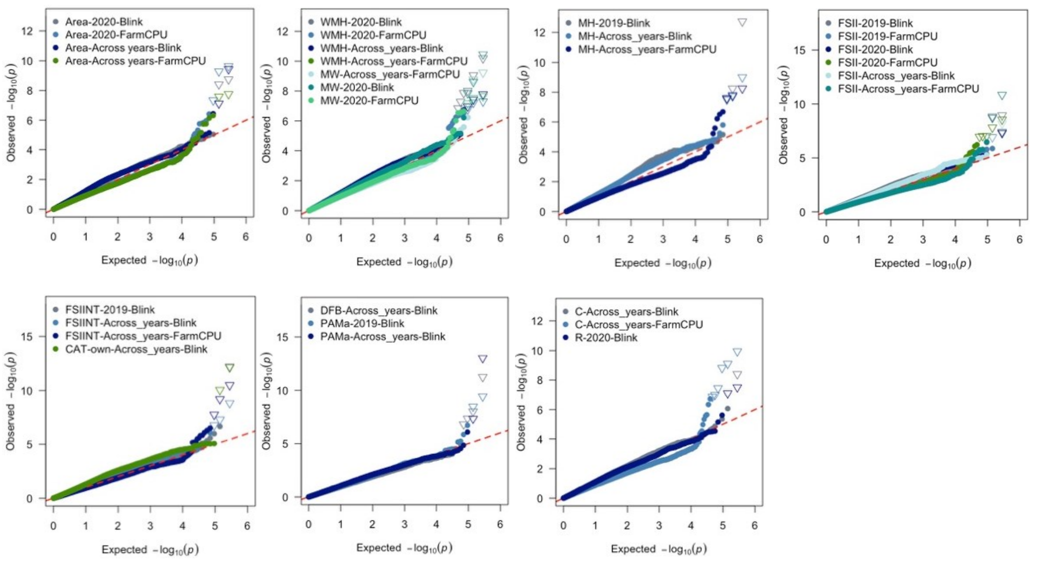


**Supplementary Figure 4**: QQplot of GWAS results on mean phenotype per year or across years and GWAS model. The legends of each plot correspond to the trait-year or across years-model.

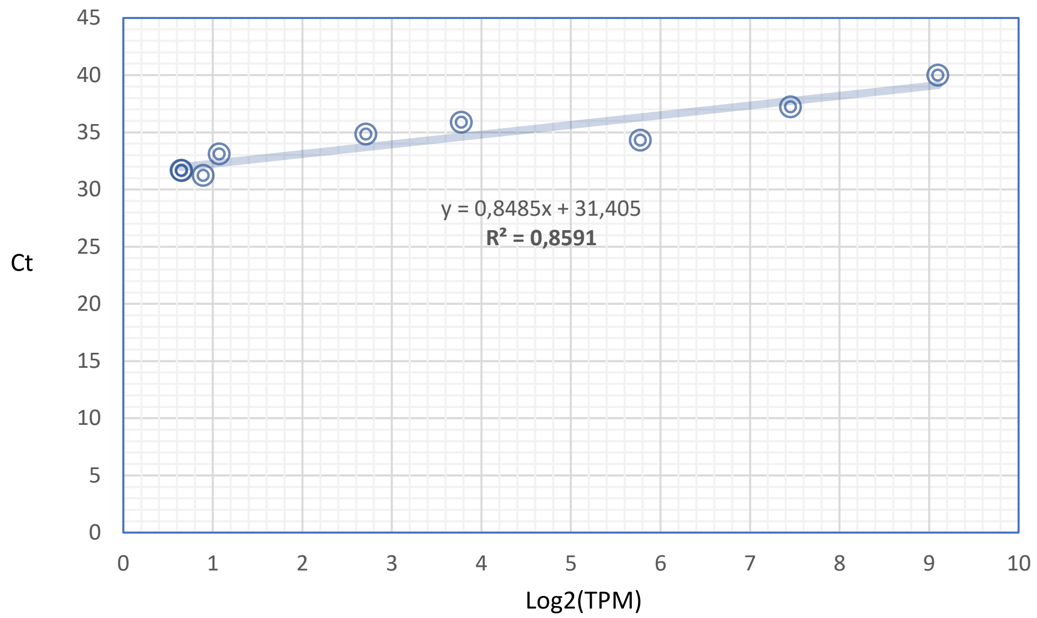


**Supplementary Figure 5**: Validation of RNA-seq data by qPCR of the HF43536 gene. The trend line is represented by the equation below and R-squared. Log2(TPM) on the X-axis and Ct on the Y-axis.

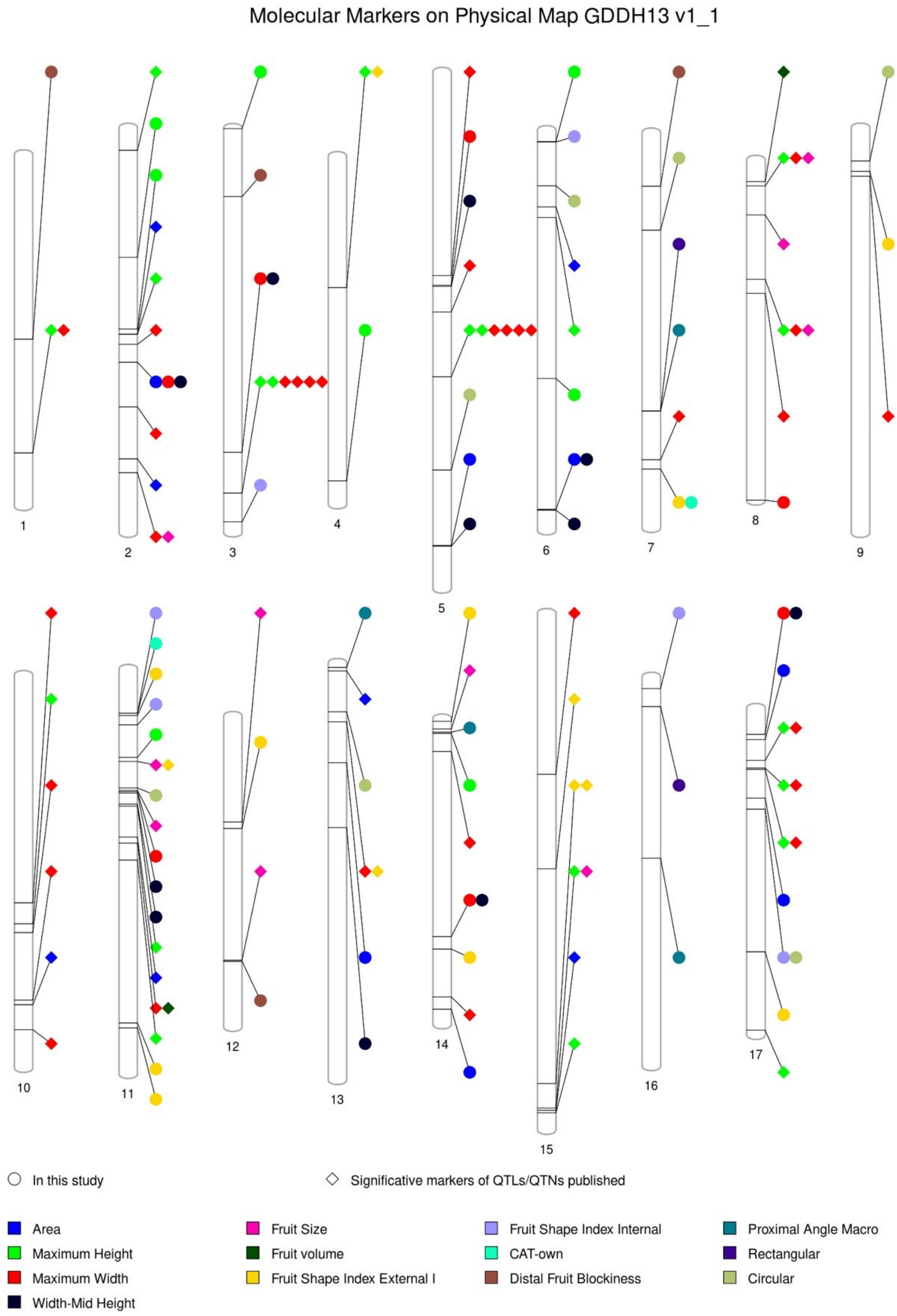


**Supplementary Figure 6**: Molecular Markers on Physical map GDDH13v1.1. Significant markers for mapping in apple fruit measures, including markers published described in some QTLs/QTNs analysis and QTNs for this study. Symbols: circle, correspond to "in this study" and diagonal, "significative markers of QTLs/QTNs published". Each color corresponds to different trait for size and shape. See more details in **Supplementary Table 7**.
